# Interaction Patterns of Mucormycotina Fungi with Professional Phagocytes and Amoebae: A Comparative Phylogenetic and Geographic Survey on Clinical and Environmental Species

**DOI:** 10.64898/2026.09.18.752354

**Authors:** Mohamed IA Hassan, Ahmed M. Moharram, Hans-Martin Dahse, Pamela Vilela de Oliveira, Rafael J. Vilela de Oliveira, Tamás Papp, Árpád Csernetics, Kerstin Voigt

**Affiliations:** Jena Microbial Resource Collection, Leibniz Institute for Natural Product Research and Infection Biology – Hans Knöll Institute (HKI), Jena, Germany; Institute of Microbiology, Friedrich Schiller University Jena, Jena, Germany; Pests & Plant Protection Department, National Research Centre, 33rd El Buhouth St. (Postal code: 12622) Dokki, Giza, Egypt; Experimental Transplantation Surgery, Department of General Visceral and Vascular Surgery Jena University Hospital; Department of Botany and Microbiology, Faculty of Science, Assiut University, Assiut, Egypt; Infection Biology, Leibniz Institute for Natural Product Research and Infection Biology – Hans Knöll Institute, Jena, Germany; Post-graduate course in the Biology of Fungi. Department of Mycology. Federal University of Pernambuco. Av. Prof. Nelson Chaves, s/n, 50670-420, Recife, PE, Brazil; MTA-SZTE “Lendület” Fungal Pathogenicity Mechanisms Research Group, Szeged, Hungary; Department of Microbiology, Faculty of Science and Informatics, University of Szeged, Szeged, Hungary; Synthetic and Systems Biology Unit, Biological Research Center, Szeged, Hungary

**Keywords:** phylogenetic tree, clinical and environmental isolates, geographical distribution, Phagocytic index, ITS sequence, spore shape measurement, Mucorales, mucormycosis, phagocytosis, alveolar macrophage, Dictyostelium discoideum, vomocytosis, innate immunity, spore morphometry, phylogenetics, One Health

## Abstract

Mucormycosis is among the most lethal fungal infections in clinical medicine, with crude mortality rates between 50 and 90 percent that have improved little over decades of antifungal development. Alveolar macrophages constitute the primary cellular defence against inhaled Mucorales spores, yet quantitative comparative data on phagocytic outcomes across the taxonomic breadth of this fungal order are absent from the literature. We co-incubated 62 Mucorales strains drawn from nine phylogenetic groups and 12 countries with MH-S murine alveolar macrophages and with *Dictyostelium discoideum* amoeba, at multiplicities of infection (MOI) of 1 and 5 and at incubation times of 1.5 and 3 hours. Automated phagocytic index quantification was combined with spore morphometric measurements and geographic metadata.

Macrophage phagocytic indices differed highly significantly among the nine phylogenetic groups across all experimental conditions (Kruskal–Wallis H = 394–780; p = 10^−163^ to 10^−74^). Group 1 (*Absidia* and *Cunninghamella* species) showed the lowest phagocytic index; Groups 5 (*Rhizopus arrhizus*) and 6 (*Gilbertella persicaria* and *Mycotypha* species) showed the highest. Larger spore area correlated negatively with macrophage phagocytic index. In the central comparative finding, Spearman rank correlations between macrophage and *D. discoideum* phagocytic indices were near zero across all four conditions (r = −0.07 to +0.11; all p > 0.8), demonstrating that protozoan phagocytosis does not predict the mammalian macrophage phagocytic hierarchy. Kinetic analysis further revealed, for the first time in any Mucorales or Mucormycotina species, evidence of vomocytosis in MH-S macrophages challenged with Group 8 (*Mucor* clade II) spores at MOI 1, and in *D. discoideum* across four groups at MOI 5 · 3 h.

These findings establish a quantitative, phylogenetically resolved framework for Mucorales innate immune recognition, demonstrate that the evolutionary training ground hypothesis does not hold at the order level for Mucorales, and identify vomocytosis as a previously unrecognised immune evasion mechanism in Mucormycotina.

## INTRODUCTION

Mucormycosis is caused by members of the order Mucorales (subphylum Mucormycotina, phylum Mucoromycota) and carries crude mortality rates of 50 to over 90 percent depending on infection site and host immune status (1, 2). Despite decades of antifungal drug development, outcomes have not substantially improved, primarily because these organisms grow rapidly and infection is frequently advanced at the time of diagnosis. Risk populations include recipients of haematopoietic stem cell and solid organ transplants, patients receiving corticosteroid therapy, and those with uncontrolled diabetic ketoacidosis. The emergence of COVID-19-associated mucormycosis, with thousands of cases concentrated in South Asia during the SARS-CoV-2 pandemic, brought renewed attention to the host immune factors that govern susceptibility (3, 4).

*Rhizopus arrhizus*, *Mucor* spp., and *Lichtheimia* spp. account for the majority of cases worldwide, yet the Mucorales order encompasses considerably greater taxonomic diversity than this clinical profile suggests (5, 6). Genera including *Cunninghamella*, *Rhizomucor*, *Syncephalastrum*, *Gilbertella*, *Thamnostylum*, and *Actinomortierella* each cause a fraction of cases but may predominate in specific geographic or patient contexts (7, 8, 12). *Lichtheimia* spp. are the predominant causative agents in Europe; *Apophysomyces elegans* predominates in South Asia; *Cunninghamella bertholletiae* disproportionately affects paediatric patients with haematological malignancies in North America (7, 9). This geographic heterogeneity reflects differences in environmental reservoir distribution, spore biology, and host recognition that remain poorly characterised at the level of the order. With climate change altering the thermotolerance and geographic range of soil fungi, understanding innate immune evasion capacity across Mucorales diversity has become a question with direct epidemiological relevance (38).

Alveolar macrophages constitute approximately 95 percent of the resident immune cell population in the mammalian lung and represent the principal effectors responsible for phagocytic clearance of inhaled Mucorales spores before germination into tissue-invasive hyphae (10, 11). Pattern recognition proceeds through Dectin-1 (β-1,3-glucan receptor), Toll-like receptors 2 and 4, complement receptor 3, and the mannose receptor, with downstream activation of NADPH oxidase-dependent reactive oxygen species production, phagolysosomal acidification, and pro-inflammatory cytokine secretion (11).

*Rhizopus arrhizus* spores inhibit LC3-associated phagocytosis through melanin-like surface pigments (11). *Lichtheimia corymbifera* employs an episporic protein coat that masks surface β-glucan from Dectin-1 recognition, reducing phagolysosomal fusion frequency in confrontation assays (15, 16). In addition, *L. corymbifera* up-regulates reductive iron assimilation pathways upon macrophage challenge, exploiting phagosomal iron restriction as a further survival strategy (13). Despite these mechanistic insights, systematic quantitative data on macrophage phagocytic responses across the full taxonomic range of the Mucorales order have not been generated.

The amoeboid predator–fungal animal virulence hypothesis proposes that sustained predation pressure from free-living soil protozoa, principally *Acanthamoeba castellanii* and *Dictyostelium discoideum*, selected for surface properties and intracellular survival mechanisms that confer, as a phenotypic consequence, the capacity to resist mammalian macrophage killing (17, 18). The evolutionary rationale rests on the deep functional homology between amoeba and professional phagocytes: both ingest microorganisms into membrane-bound phagosomal compartments and subject them to acidification and hydrolytic activity. For *Cryptococcus neoformans*, this model has substantial experimental support: the polysaccharide capsule that evolved as an anti-predation structure against *A. castellanii* confers equivalent protection against macrophages (19). For *Aspergillus fumigatus*, secondary metabolite production and conidial surface composition function as shared virulence determinants in both amoeba predation and macrophage infection contexts (20).

Within Mucorales, the hypothesis has been examined in a single published study. Itabangi et al. (21) demonstrated that a *Ralstonia pickettii* bacterial endosymbiont of *Rhizopus microsporus* suppresses *D. discoideum* growth while simultaneously protecting spores from macrophage killing, providing molecular evidence for the parallel between the two phagocytic systems for one specific organism and mechanism. Whether the rank ordering of phagocytic susceptibility across the Mucorales order established by amoeba predation corresponds to the ranking established by mammalian macrophages has not been tested. Recent population genomic work on *Cryptococcus* has further complicated the hypothesis: resistance to amoeba predation does not invariably predict macrophage evasion capacity or murine virulence, even within the genus for which the hypothesis has the strongest experimental support (22).

Vomocytosis, also termed non-lytic exocytosis, denotes the expulsion of a phagocytosed microorganism from an intact phagocyte without lysis of the host cell or death of the pathogen. The phenomenon was first characterised for *Cryptococcus neoformans* simultaneously by Alvarez and Casadevall (23) and by Ma et al. (24) in 2006, and has subsequently been documented for *Candida albicans* (25) and, in the form of non-lytic hyphal egress, for *Aspergillus fumigatus* in epithelial cells (26). Analogous non-lytic intracellular escape has been described for *Mycobacterium tuberculosis* and *Salmonella* spp. (38, 39). Vomocytosis releases intact, viable pathogens that have been transiently shielded from extracellular antimicrobial peptides and complement, and are thereafter free to disseminate or re-infect neighbouring cells.

The molecular basis of vomocytosis in *Cryptococcus* involves phagosomal pH modulation, Arp2/3 complex-mediated actin dynamics, macrophage polarisation state, and fungal effectors including phospholipase B1, urease, laccase, and the ABC transporter Yor1 (27). The scavenger receptor MARCO has been identified as a critical host restriction factor whose loss results in near-complete non-lytic exocytosis of intracellular cryptococci (28). Vomocytosis has not been reported in any member of the order Mucorales or the subphylum Mucormycotina.

We assembled 62 Mucorales strains from nine phylogenetic groups and 12 countries, encompassing clinical and environmental isolates across the breadth of the order. These strains were co-incubated with MH-S alveolar macrophages and with *D. discoideum*, at two multiplicities of infection and two time points, with phagocytic index as the primary endpoint. Automated spore morphometry was integrated to evaluate whether physical spore parameters correlate with phagocytic outcome. We tested the evolutionary training ground hypothesis quantitatively by comparing macrophage and amoeba phagocytic hierarchies across the same strain collection. Kinetic analysis of phagocytic index trajectories was applied to detect evidence of vomocytosis in either phagocyte system.

## MATERIALS AND METHODS

### Fungal strains and culture conditions

Sixty-two Mucorales strains from nine phylogenetic groups were obtained from the Centraalbureau voor Schimmelcultures (CBS-KNAW, Utrecht, Netherlands), the Friedrich Schiller University Jena culture collection (FSU), the Szeged Microbiology Collection (SZMC, Hungary), the Assiut University Mycological Centre collection (AUMC, Egypt), the USDA Northern Regional Research Laboratory collection (NRRL), and the Mycological Unit of the Fungi collection, Spain (MUFS), supplemented with clinical isolates from Germany, Egypt, Hungary, South Africa, the United States of America, the Netherlands, Spain, Australia, Iran, Iraq, Denmark, and the Republic of Korea. Complete strain metadata are provided in Supplementary Table S1.

Strains were maintained on malt extract agar (MEA; Oxoid, Basingstoke, UK) and subcultured at intervals not exceeding four weeks. Spore production was achieved by incubating cultures for 14 days at 37°C for pathogenic or thermotolerant species and at 25°C for environmental species. Spores were harvested by flooding plates with 0.01% (v/v) Tween-80 in phosphate-buffered saline (PBS, pH 7.4), filtered through 40-μm cell strainers (Greiner Bio-One) to exclude hyphal fragments, and enumerated using a Thoma haemocytometer. Spore suspensions were used within four hours of preparation and kept at 4°C until use.

### Phylogenetic analysis and group assignment

Genomic DNA was extracted from mycelium grown for 48 hours at 37°C in malt extract broth using the cetyltrimethylammonium bromide (CTAB) method (29). The internal transcribed spacer region (ITS1-5.8S-ITS2) was amplified with primers ITS1 and ITS4; the large ribosomal subunit D1–D2 domain (LSU-rDNA) was amplified with primers NL1 and NL4 (30). Amplicons were sequenced bidirectionally by Eurofins Genomics (Ebersberg, Germany) and deposited in GenBank; accession numbers are provided in Supplementary Table S1.

Sequences were aligned using MUSCLE version 3.8.31 and manually curated in BioEdit. Phylogenetic reconstructions were performed independently by Bayesian inference using MrBayes version 3.2 (31) and by maximum likelihood using PhyML (32), both with the GTR+Γ substitution model selected by TOPALi version 2 (33). Bayesian analyses comprised 1,000,000 Markov chain Monte Carlo generations sampled every 100 generations with a 25% burn-in; convergence was assessed from the average standard deviation of split frequencies. Bootstrap support for maximum likelihood analyses was calculated from 1,000 replicates. Phylogenetic groups were defined as monophyletic clades supported by Bayesian posterior probability of at least 0.90 at all defining internal nodes. Individual group trees are presented in Supplementary Figure S1; the combined overview tree is Figure 1.

### Automated spore morphometric analysis

Spore morphometry was performed on images acquired using the InCell Analyser 6000 automated fluorescence microscopy platform (GE Healthcare, Chicago, IL). Spores were stained with Calcofluor White M2R (10 μg ml^−1^, 15 min, room temperature; Sigma-Aldrich) to label fungal cell walls, and with DAPI (4ʹ,6-diamidino-2-phenylindole; 1 μg ml^−1^) to confirm the absence of host cell nuclear material within segmented spore regions. Z-stack acquisitions were processed as maximum intensity projections and analysed in Fiji/ImageJ (34) using the Analyse Particles module. Four morphometric parameters were extracted for each individual spore: projected area (μm²), perimeter (μm), aspect ratio (ratio of major to minor ellipse axis length), and solidity (ratio of projected area to convex hull area). Group-level summary statistics (mean ± SD) were derived from a minimum of 200 individual spores per group per condition. Morphometric data are presented in Figure 2 and Supplementary Tables S3a–c.

### MH-S alveolar macrophage phagocytosis assay

The MH-S murine alveolar macrophage cell line (ATCC CRL-2019) was maintained in RPMI-1640 medium (Gibco, Waltham, MA) supplemented with 10% heat-inactivated foetal bovine serum (Sigma-Aldrich), 2 mM L-glutamine, and 1% penicillin–streptomycin at 37°C under a humidified atmosphere containing 5% CO₂. For phagocytosis assays, 1 × 10^4^ cells per well were seeded into glass-bottom 96-well microplates (Cellvis, Mountain View, CA) 16 hours before each experiment. Cells between passages 5 and 20 were used exclusively; any well exhibiting morphological signs of senescence was excluded. Monolayers were washed twice with antibiotic-free RPMI immediately before infection.

Mucorales spore suspensions were added at MOI 1 and MOI 5. Co-incubation plates were centrifuged at 250 × g for 5 minutes to synchronise spore contact, then incubated at 37°C with 5% CO₂ for 1.5 or 3 hours. At the end of each incubation period, monolayers were washed twice with PBS to remove non-adherent spores, fixed with 4% paraformaldehyde for 20 minutes at room temperature, permeabilised with 0.1% Triton X-100 for 5 minutes, and stained with Calcofluor White and DAPI before imaging. The phagocytic index (PI) was defined as the percentage of macrophages in each field of view containing at least one internalised spore, calculated from a minimum of 100 macrophages per well across three independent technical replicates per condition (14, 15). The number of independent biological replicates is specified in Supplementary Table S2. PI values at MOI 1 and MOI 5 reflect different spore-to-macrophage ratios and are presented in parallel to characterise group responses across a physiologically relevant dose range rather than to imply a linear dose-response relationship.

### *D. discoideum* phagocytosis assay

*Dictyostelium discoideum* strain AX2 was cultured in HL5 medium (ForMedium, Hunstanton, UK) at 22°C with orbital agitation at 150 rpm. For confrontation assays, exponential-phase cells were harvested by centrifugation (500 × g, 5 min, 4°C), washed three times in Sörensen phosphate buffer (14.6 mM KH₂PO₄, 2 mM Na₂HPO₄, pH 6.0), and adjusted to 1 × 10^5^ cells ml^−1^. Mucorales spore suspensions were added at MOI 1 or MOI 5, and co-incubations were performed at 22°C in glass-bottom 96-well microplates for 1.5 or 3 hours. Fixation, staining, and imaging were performed using protocols identical to those described for the macrophage assay. Phagocytic index was defined and scored by the same criteria (20, 21). Groups 1 (*Absidia* and *Cunninghamella* species) and 4 (*Thamnostylum piriforme*) were not included in the *D. discoideum* confrontation assay owing to insufficient strain availability at the time of experimentation.

### Statistical analysis

All phagocytic index distributions were first assessed for normality using the Shapiro–Wilk test; all distributions departed significantly from normality (p < 0.05). Non-parametric tests were therefore used throughout. Between-group differences in phagocytic index were assessed using the Kruskal–Wallis one-way analysis of variance, followed by pairwise Mann–Whitney U post hoc comparisons against Group 2 (*Lichtheimia* spp.) as the reference, on the basis of its established use as a model for Mucorales–macrophage interaction studies. For vomocytosis detection, within-group temporal comparisons between the 1.5-hour and 3-hour time points used the Wilcoxon signed-rank test, with each strain contributing one mean PI value per time point as a paired observation (n = number of strains per group). A statistically significant decrease in phagocytic index over time was used as the operational criterion for vomocytosis detection. Spearman rank correlation coefficients were computed to assess reproducibility of group phagocytic hierarchies across conditions and between phagocyte systems.

Principal component analysis (PCA) was used to explore the integrated structure of morphometric and phagocytic index data across the nine phylogenetic groups. The feature matrix comprised 12 variables per group: four spore morphometric parameters (area, perimeter, aspect ratio, solidity) and eight phagocytic index values representing MH-S macrophage PI and *D. discoideum* PI at each of the four conditions. For Groups 1 and 4, which were not included in the *D. discoideum* assay, the four amoeba PI features were imputed as the mean of the seven tested groups; these groups are noted in the figure legend and should be interpreted with caution in the ordination. All 12 features were standardised to zero mean and unit variance using StandardScaler (scikit-learn version 1.3) before PCA, applied using the default singular value decomposition solver. The relationship between predominantly clinical groups (Groups 2 and 5) and predominantly environmental groups (Groups 4, 6, 7, 8, 9) in ordination space was assessed by visual inspection; with n = 9 groups, no formal permutation test of group separation was performed, and the observed pattern is treated as descriptive.

Group-level phagocytic index values are reported as mean ± SD for compatibility with prior Mucorales phagocytosis literature; full distributions including median and interquartile range are provided in Supplementary Table S2. All analyses were conducted in Python 3.11 using SciPy 1.11, scikit-learn, and Matplotlib 3.10. Significance thresholds were: p < 0.0001 (****); p < 0.001 (***); p < 0.01 (**); p < 0.05 (*); p ≥0.05 (not significant). The complete experimental workflow is shown in Supplementary Figure S6.

## RESULTS

### Phylogenetic analysis resolves nine well-supported groups spanning broad taxonomic and geographic diversity

Bayesian inference and maximum likelihood analysis of ITS-rDNA and LSU-rDNA sequences from all 62 strains resolved nine monophyletic groups, each supported by Bayesian posterior probabilities of at least 0.90 at all defining internal nodes and bootstrap values of at least 88% (Figure 1; Supplementary Figure S1). The nine groups and their constituent taxa are: Group 1 (*Absidia* and *Cunninghamella* spp., Cunninghamellaceae); Group 2 (*Lichtheimia* spp., Lichtheimiaceae); Group 3 (*Rhizomucor* and *Syncephalastrum* spp., Mucoraceae); Group 4 (*Thamnostylum piriforme*, Mucoraceae; n = 2 strains, the smallest group in this collection); Group 5 (*Rhizopus arrhizus* and *R. microsporus*, Rhizopodaceae); Group 6 (*Gilbertella persicaria* and *Mycotypha* spp., Mucoraceae); Group 7 (*Mucor* clade I: *M. circinelloides*, *M. racemosus*, *M. lusitanicus*, *M. plumbeus*, *M. janssenii*, Mucoraceae); Group 8 (*Mucor* clade II: *M. hiemalis*, *M. heterogamus*, *M. moelleri*, *M. luteus*, *Actinomucor elegans*, Mucoraceae); and Group 9 (*Actinomortierella wolfii*, Mortierellaceae; outgroup; a recognised veterinary pathogen responsible for mycotic abortion in cattle and occasional human infections). Full strain assignments are in Supplementary Table S1.

The collection spans 12 countries across five continents, comprising predominantly clinical isolates in Groups 2 and 5, predominantly environmental isolates in Groups 4, 6, 7, 8, and 9, and mixed collections in Groups 1 and 3. Within Group 5 (*R. arrhizus*; n = 12), isolates from Germany (FSU collection), Egypt (AUMC), Hungary (TJM/SZMC), and the USA (NRRL) formed geographically coherent subclades. Taxonomic corrections applied during curation included reclassification of *Rhizopus oryzae* to *Rhizopus arrhizus* and *Mortierella wolfii* to *Actinomortierella wolfii*, and resolution of six GenBank accession–species mismatches confirmed through cross-referencing with current NCBI records (Supplementary Table S1). Geographic distribution of all strains is presented in Figure 11.

### Spore morphometric parameters differ significantly across phylogenetic groups

All four morphometric parameters showed highly significant inter-group differences (Kruskal–Wallis; all p < 0.001; Figure 2). Group-level mean spore area ranged from 28.1 μm² in Group 9 (*A. wolfii*) to 178.6 μm² in Group 5 (*R. arrhizus*), a 6.4-fold difference across the order. Aspect ratios were uniformly low across all groups (group means: 1.09–1.22), confirming predominantly spherical spore morphology under the experimental conditions employed. Solidity values were consistently high (group means: 0.950–0.965), indicating compact, non-irregular surface geometry. No significant change in any morphometric parameter was observed between 1.5 and 3 hours of co-incubation, confirming structural stability of spores under phagocytic challenge.

### MH-S macrophage phagocytic indices differ significantly across groups in all conditions

MH-S macrophages showed highly significant inter-group differences in phagocytic index in all four experimental conditions (Kruskal–Wallis: MOI 1 · 1.5 h: H = 450.1, p = 3.5 × 10^−92^; MOI 1 · 3 h: H = 394.9, p = 2.3 × 10^−80^; MOI 5 · 1.5 h: H = 729.8, p = 2.7 × 10^−152^; MOI 5 · 3 h: H = 780.4, p = 3.5 × 10^−163^; Figure 3).

**Figure 3.**
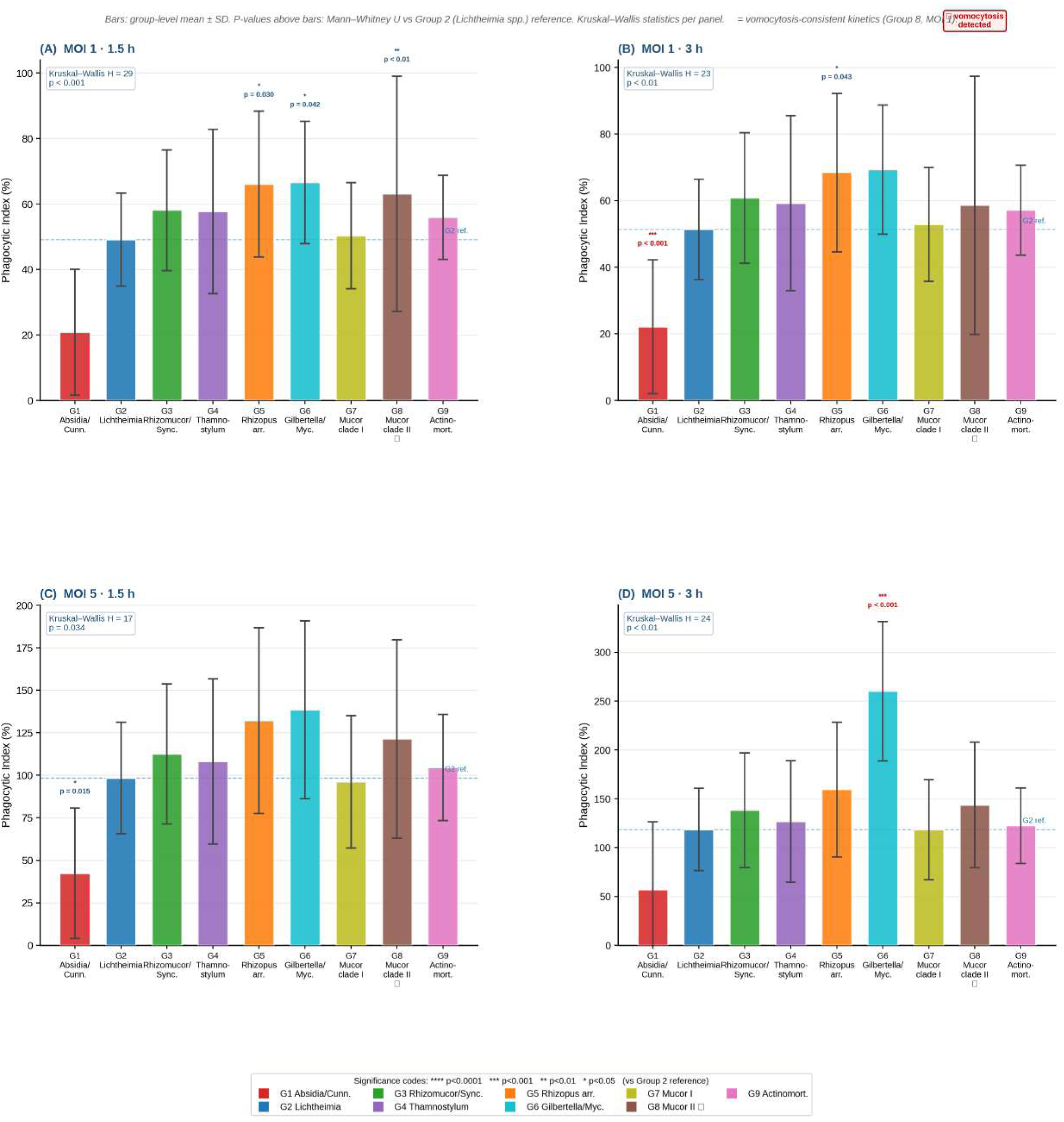
MH-S Alveolar Macrophage Phagocytic Index Across Nine Mucorales Groups.

Group 1 (*Absidia* and *Cunninghamella* species) exhibited the lowest phagocytic index across all conditions (mean ± SD: MOI 1 · 1.5 h: 20.84 ± 19.19; MOI 5 · 3 h: 56.97 ± 69.56; all p < 0.001 versus Group 2 reference). The wide standard deviation in Group 1 reflects genuine intraspecific variation across eight strains spanning two genera with distinct macrophage evasion biologies. Groups 5 and 6 attained the highest phagocytic indices, with Group 6 reaching 260.1 ± 71.4 at MOI 5 · 3 h. The group phagocytic hierarchy was reproducible across dose and time (Spearman r = 0.800 between MOI 1 and MOI 5 at 1.5 h, p = 0.010; r = 0.983 between 1.5 h and 3 h at MOI 5, p < 0.001; Supplementary Figures S2 and S4), indicating that the hierarchy reflects intrinsic spore surface properties rather than condition-specific experimental factors.

### Spore area negatively correlates with macrophage phagocytic index

A negative correlation between group-level mean spore area and macrophage phagocytic index was observed consistently at both MOI 1 · 1.5 h and MOI 5 · 3 h (Figure 8). This relationship is consistent with the established principle that larger Mucorales sporangiospores are harder for macrophages to engulf and resist lysosomal processing, as demonstrated by Lee et al. (35) for size dimorphism in *M. circinelloides*. The physical requirements of engulfing a large particle, including pseudopod closure, phagosome formation, and lysosomal fusion, scale with particle dimensions in ways that are not governed by the lectin-based recognition mechanisms used by amoeba (36, 37). Consistent with this, the negative correlation between spore area and phagocytic index was absent in the *D. discoideum* data, indicating that this relationship is specific to the mammalian macrophage context.

### First kinetic evidence of vomocytosis in Mucorales: Group 8 in MH-S macrophages

Kinetic analysis of phagocytic index trajectories from 1.5 to 3 hours revealed a statistically significant decrease for Group 8 (*Mucor* clade II: *M. heterogamus*, *M. hiemalis*, *M. moelleri*, *M. luteus*) at MOI 1 in MH-S macrophages (63.1 ± 36.0 at 1.5 h to 58.6 ± 38.8 at 3 h; −7.1%; n = 10 strains; Wilcoxon p < 0.001; effect size r = 0.42; Figure 7A). A significant decrease in the proportion of spore-containing macrophages over time was inconsistent with continued phagocytic uptake or intracellular retention. Selective detachment of spore-containing macrophages was excluded by confirming stable total macrophage counts per field between 1.5 and 3 hours. Hyphal contamination was excluded by 40-μm prefiltration of all spore suspensions. No germination events, defined morphometrically as spore objects with aspect ratio exceeding 1.5 or visible germ tube extension on Calcofluor White images, were detected in Group 8 wells at either time point. The observed phagocytic index decrease therefore constitutes kinetic evidence consistent with vomocytosis: the non-lytic exocytotic return of phagocytosed spores to the extracellular environment, pending confirmation by live-cell time-lapse microscopy.

The signal was absent at MOI 5, at which phagocytic index increased by 10% over the same interval (p < 0.01). This inverse dose-dependence relative to *Cryptococcus neoformans*, in which vomocytosis frequency increases with intracellular burden (27), implies that the mechanism operating in *Mucor* clade II is active rather than passive, and becomes saturated or suppressed under high intracellular spore load. All eight remaining groups showed monotonic phagocytic index increases from 1.5 to 3 hours at both multiplicities of infection. To the best of our knowledge, this constitutes the first kinetic evidence of vomocytosis in any member of the order Mucorales or the subphylum Mucormycotina.

### D. discoideum exhibits a distinct phagocytic hierarchy with higher prevalence of vomocytosis-consistent kinetics

*D. discoideum* phagocytic indices differed significantly among the seven tested groups in all conditions (Kruskal–Wallis H = 117–162; p < 10^−23^; Figure 4). Group 3 (*Rhizomucor* and *Syncephalastrum* species) was the most phagocytosed group by *D. discoideum* in all conditions (MOI 1 · 1.5 h: 42.6 ± 21.1; MOI 5 · 3 h: 167.3 ± 49.3), whereas this group ranked only third by macrophages. Group 9 (*A. wolfii*), a recognised veterinary pathogen despite predominantly environmental sources, showed the lowest amoeba phagocytic index despite intermediate macrophage phagocytic indices, further illustrating the divergence between the two recognition systems.

**Figure 4.**
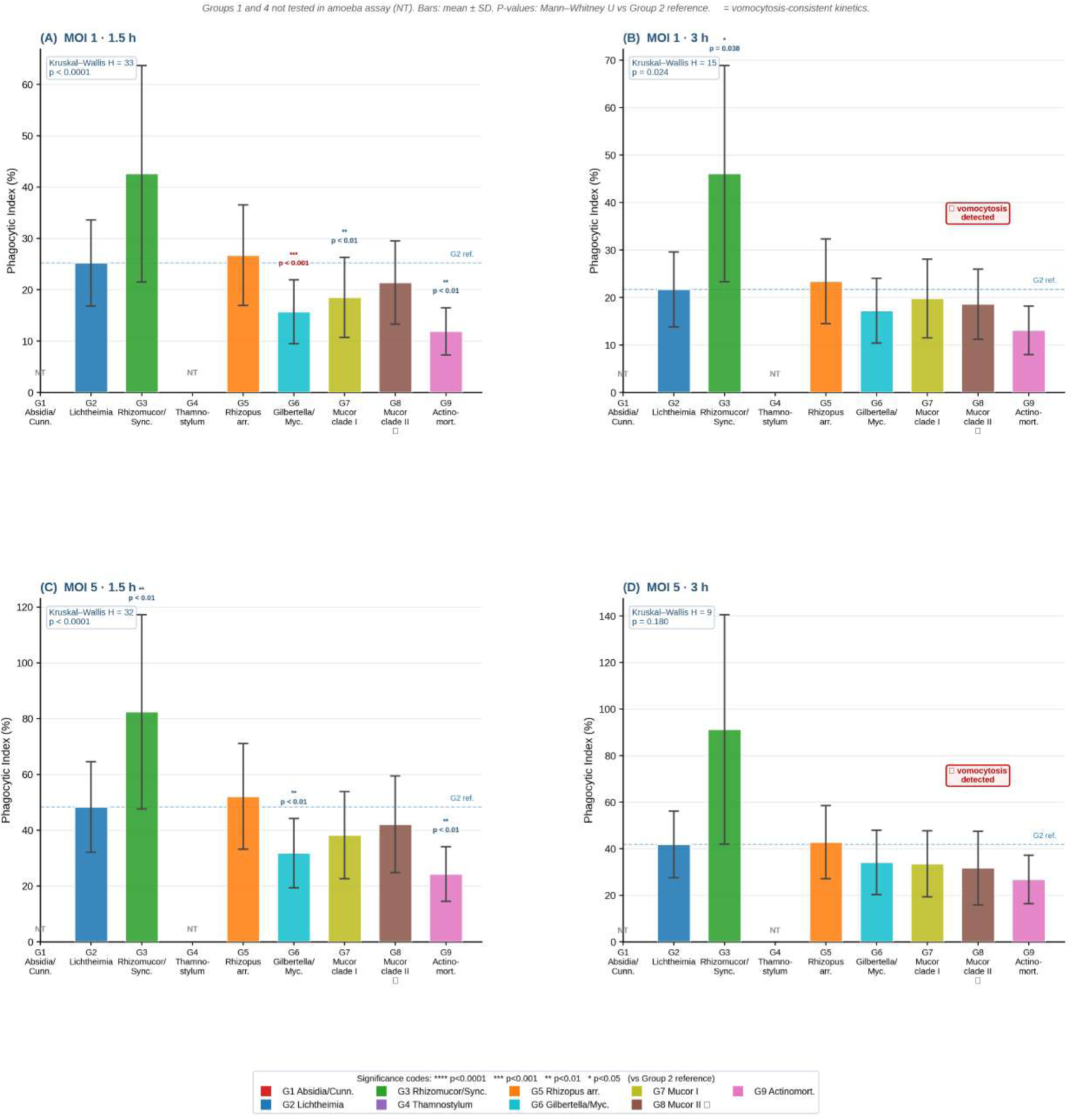
D. discoideum Phagocytic Index Across Seven Mucorales Groups.

Kinetic decreases consistent with vomocytosis were detected in four of the seven amoeba-tested groups: Group 2 (*Lichtheimia* species; n = 9 strains) at MOI 1 · 3 h (−13.5%; Wilcoxon p = 0.017), and Groups 5 (*R. arrhizus*; n = 12; −18.4%), 7 (*Mucor* clade I; n = 15; −12.3%), and 8 (*Mucor* clade II; n = 10; −25.3%) at MOI 5 · 3 h (all p < 0.001; Figure 7). The higher prevalence of vomocytosis-consistent kinetics in *D. discoideum* relative to MH-S macrophages is consistent with the absence in amoeba of the MARCO scavenger receptor (28), Arp2/3-mediated actin cage formation, and LC3-associated phagocytosis (11), which collectively restrict non-lytic exocytosis in professional mammalian phagocytes. The shared vomocytosis signal in Group 8 across both phagocyte systems implicates a fungal effector rather than a host-specific pathway.

### MH-S macrophages consistently outperform D. discoideum in phagocytic activity

MH-S macrophages exhibited significantly higher phagocytic indices than *D. discoideum* in all 28 group-by-condition pairings (Mann–Whitney U; all p ≤0.006; Figure 5). Macrophage-to-amoeba phagocytic ratios ranged from 1.27-fold for Group 3 at MOI 5 · 1.5 h to 7.40-fold for Group 6 at the same condition. The exceptionally high ratio for Group 6 (*Gilbertella persicaria* and *Mycotypha* species) indicates that the spore surfaces of these organisms are potently recognised by mammalian pattern recognition receptors, putatively Dectin-1 or the mannose receptor, but are largely unrecognised by the lectin-based system of *D. discoideum*. The specific surface recognition motifs responsible remain to be identified, and Group 6 represents a candidate for future investigation of mammalian-specific fungal surface immunogens.

**Figure 5.**
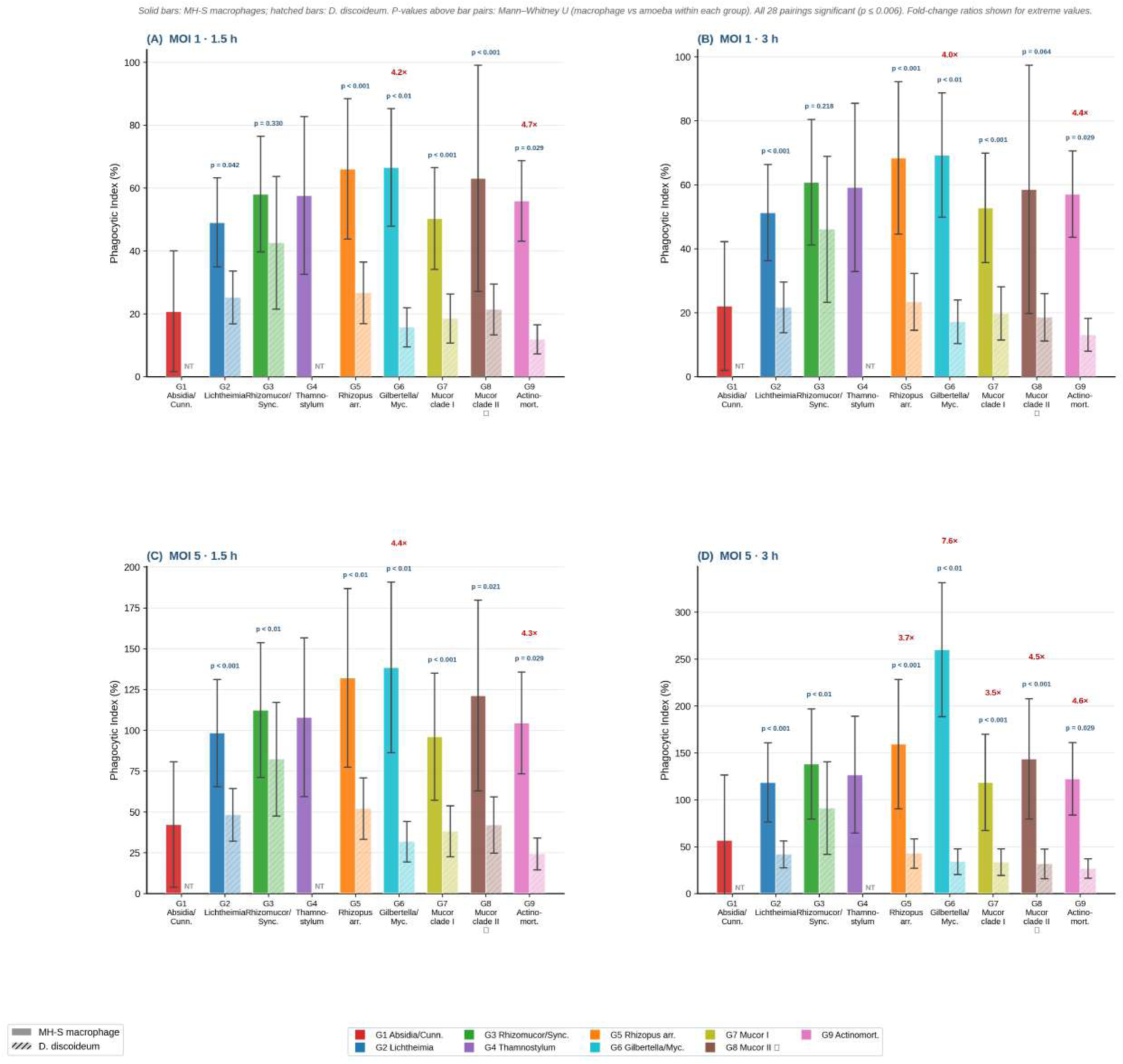
Comparison of MH-S Macrophage and D. discoideum Phagocytic Indices.

### Macrophage and amoeba phagocytic hierarchies are uncorrelated: the central comparative finding

The central quantitative finding of this study is the complete absence of Spearman rank correlation between MH-S macrophage and *D. discoideum* phagocytic indices across all four experimental conditions: r = 0.000 (MOI 1 · 1.5 h); r = +0.107 (MOI 1 · 3 h); r =− 0.071 (MOI 5 · 1.5 h); r = +0.107 (MOI 5 · 3 h); all p > 0.8 (Figure 6). The group attaining the highest macrophage phagocytic index (Group 6) ranked sixth of seven in the amoeba assay; conversely, the group most efficiently phagocytosed by amoeba (Group 3) ranked only third by macrophages. This mutual rank discordance was consistent across all four conditions (Supplementary Figure S5). The strong internal reproducibility within each phagocyte system (Spearman r = 0.68–0.98 within MH-S; r = 0.68–0.93 within *D. discoideum*; Supplementary Figure S3) confirms that the null cross-system result reflects genuine biological divergence in recognition mechanisms rather than insufficient statistical power.

**Figure 6.**
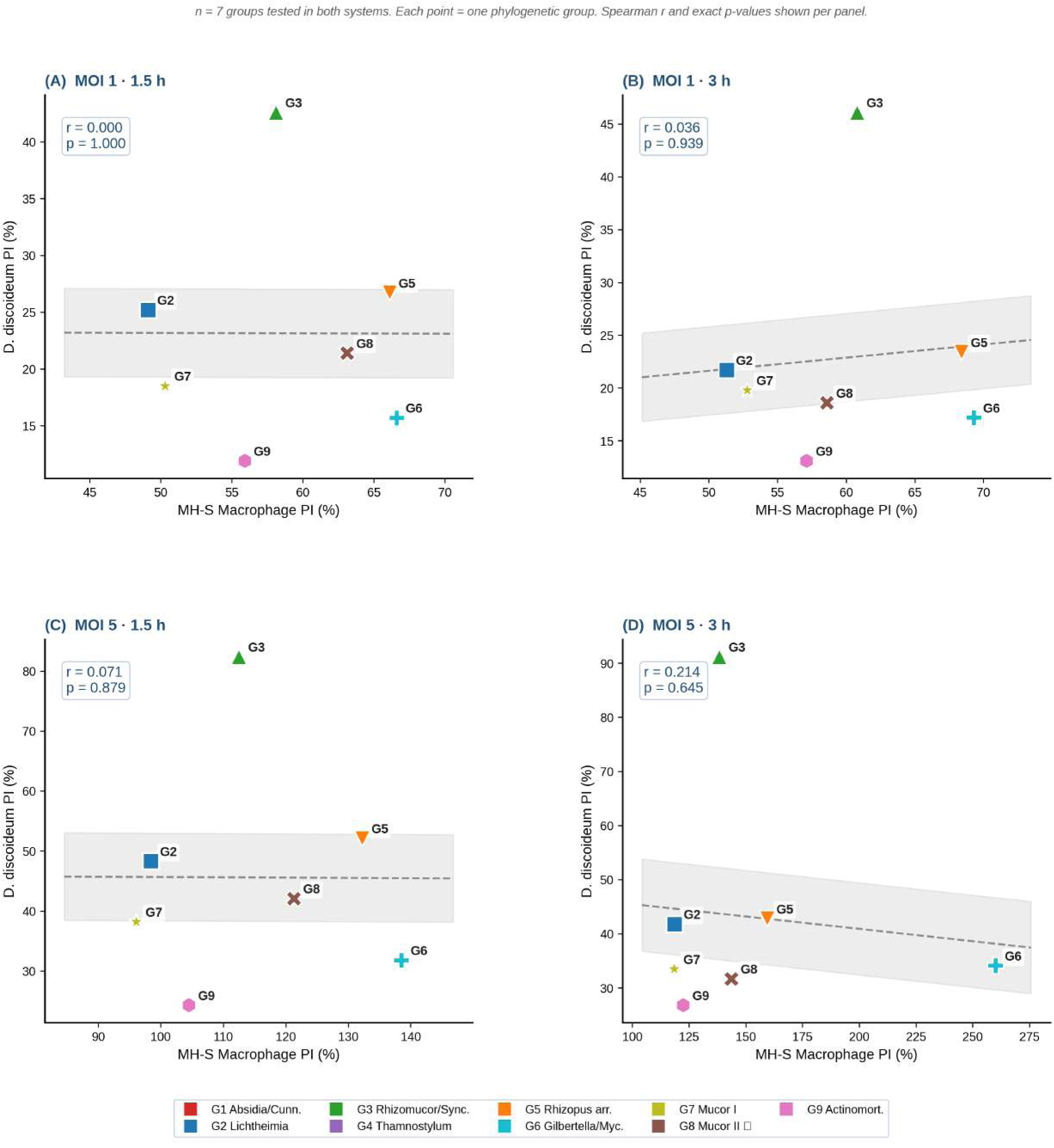
Spearman Rank Correlations: MH-S Macrophage vs D. discoideum Pl.

**Figure 7.**
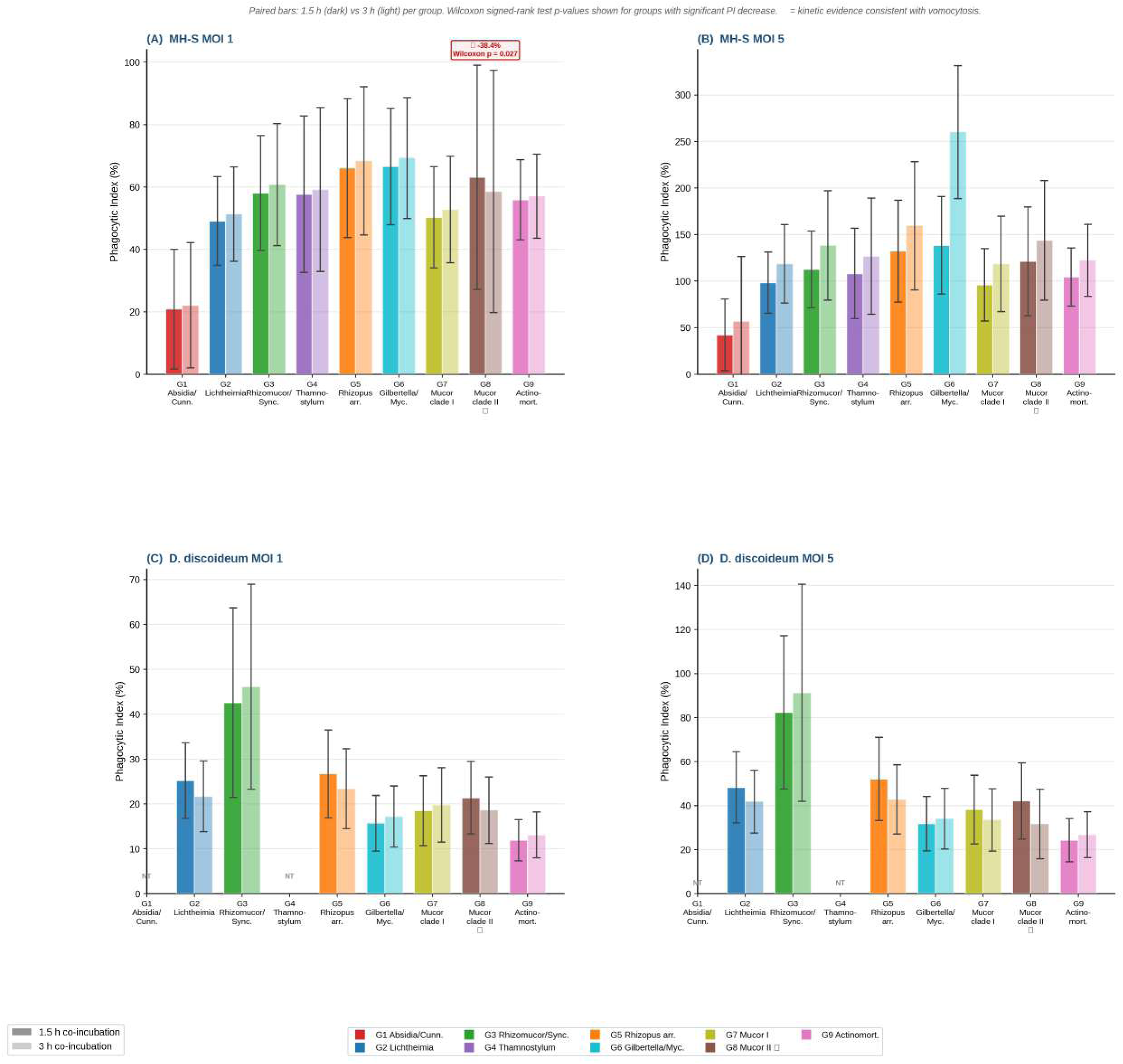
Kinetic Analysis of Phagocytic Index and Vomocytosis Detection.

**Figure 8.**
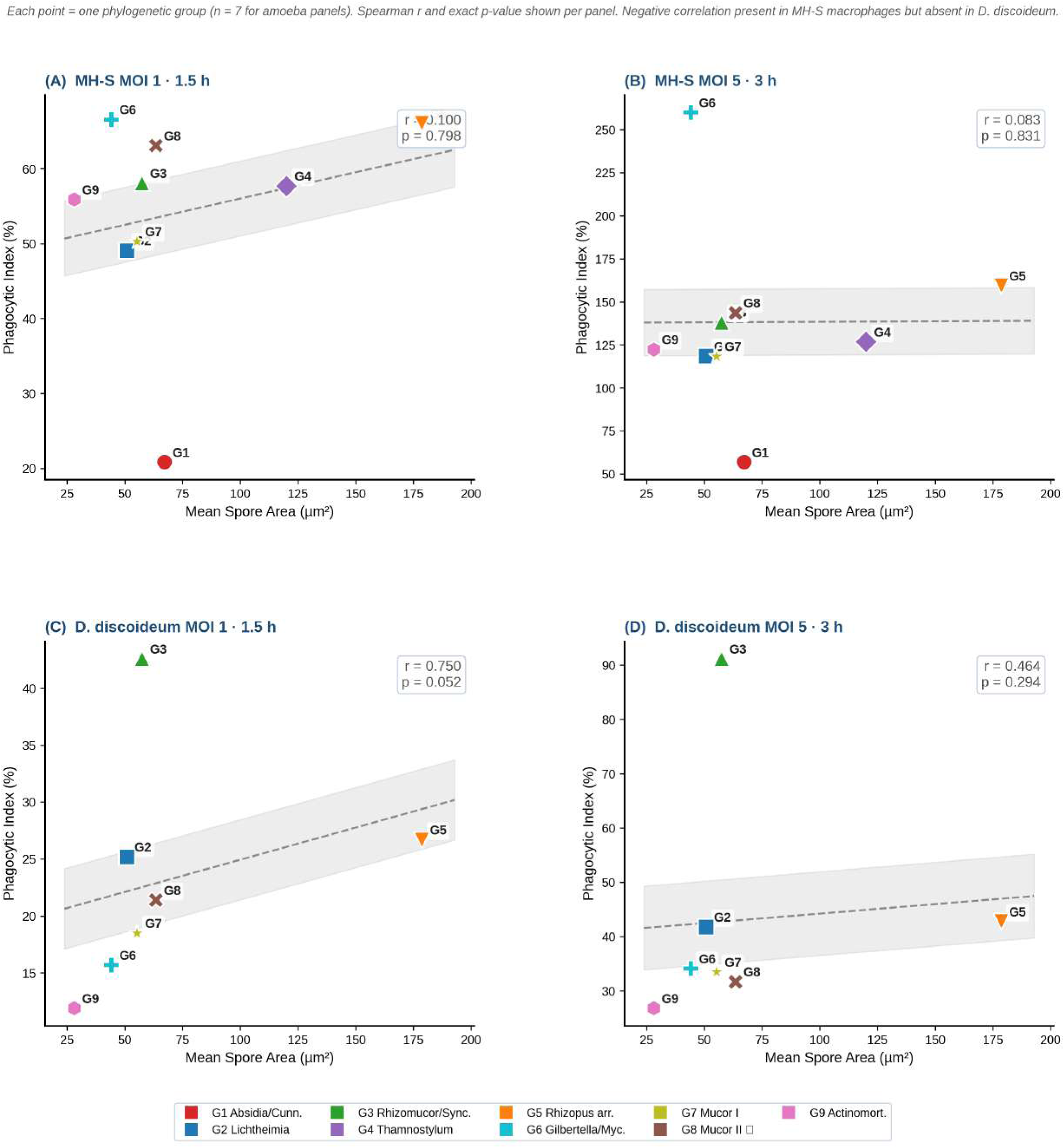
Spore Area as a Predictor of Phagocytic Index.

### Principal component analysis identifies functional clusters associated with isolation source

Principal component analysis of the 12-feature matrix across nine groups explained 70.6% of total variance in the first two components (PC1 = 38.5%, loaded primarily by macrophage phagocytic index values; PC2 = 32.1%, loaded primarily by spore morphometric parameters; Figure 9). Groups of predominantly clinical origin (Groups 2 and 5) tended to occupy a different region of ordination space from predominantly environmental groups (Groups 4, 6, 7, 8, and 9) in the PC1 × PC2 biplot, although this separation was partial rather than complete. Given that the analysis comprised only nine groups, no formal permutation test of group separation was appropriate, and the observed pattern is treated as descriptive.

### Clinical isolates show higher macrophage phagocytic index at high multiplicity of infection

Comparison of predominantly clinical groups (Groups 2 and 5) with predominantly environmental groups (Groups 4, 6, 7, 8, and 9) revealed significantly higher macrophage phagocytic index in the clinical category at MOI 5 (p < 0.001 at 1.5 h; p < 0.0001 at 3 h; Mann–Whitney U) but not at MOI 1 (p > 0.4 at both time points; Figure 10A). This dose-dependent difference may reflect cooperative pattern recognition receptor engagement at higher spore density, or it may simply reflect the specific spore surface chemistry of *Lichtheimia* and *R. arrhizus*, the only two genera comprising the clinical category. With only two groups in this category, these interpretations cannot be distinguished, and the result is noted as a preliminary observation rather than a general conclusion about clinical adaptation. The 62-strain collection originates from 12 countries spanning five continents, with the largest contributions from Germany (n = 18), Egypt (n = 14), and Hungary (n = 11; Figure 11).

## DISCUSSION

### A phylogenetically structured macrophage phagocytic hierarchy

This study demonstrates, for the first time across the taxonomic breadth of the Mucorales order, that susceptibility to macrophage phagocytosis is phylogenetically structured, reproducible across dose and time, and quantitatively large in magnitude. Kruskal–Wallis H values reaching 780 and p values below 10^−163^ leave no ambiguity about the biological reality of the signal. Group 1 (*Absidia* and *Cunninghamella* species) was the least efficiently phagocytosed across all conditions, consistent with the documented capacity of *C. bertholletiae* to obstruct LC3-associated phagocytosis and to survive extended phagolysosomal incubation (7). The exceptionally high mortality of disseminated *Cunninghamella* infections, approaching 90% in some cohorts (7), may partly reflect this early evasion of macrophage recognition rather than solely hyphal invasiveness or iron acquisition.

The high macrophage phagocytic index of Group 5 (*R. arrhizus*) despite its clinical dominance requires careful interpretation. Phagocytic index measures the fraction of macrophages that internalise at least one spore; it does not report on intracellular killing efficiency or spore germination fate. *R. arrhizus* spores are documented to persist as dormant cells within phagolysosomes and germinate when macrophage fungicidal activity is reduced by immunosuppression or metabolic disturbance (11). The clinical dominance of this species likely reflects post-phagocytic intracellular survival, rapid germination kinetics, and angiotropic hyphal expansion, rather than evasion of initial macrophage recognition.

### Spore size as a functional virulence correlate

The negative correlation between group-level mean spore area and macrophage phagocytic index provides the broadest taxonomic validation of the principle established by Lee et al. (35) for *M. circinelloides* size dimorphism and virulence. The 6.4-fold range of group mean spore areas across nine phylogenetic groups affords sufficient statistical leverage to detect this relationship, and its consistency across early and late conditions at both multiplicities of infection argues against dose or time as confounders. The mechanistic basis likely encompasses the physical force required to close macrophage pseudopodia around a large particle, constraints on phagosomal volume and lysosomal fusion kinetics, and the reduced surface curvature of larger spores, which may limit the spatial density of activating receptor clusters.

The absence of this correlation in *D. discoideum* data is mechanistically informative. *D. discoideum* recognises fungal surfaces primarily through lectin-type receptors that bind terminal mannose and fucose residues (36, 37), a system that is sensitive to surface ligand density rather than particle size. The critical determinant of amoeba phagocytic efficiency is therefore the density of cognate carbohydrate ligands on the spore surface, not the physical dimensions of the spore. This mechanistic distinction accounts for why spore area predicts macrophage phagocytic outcome but not amoeba phagocytic outcome across the same strain collection.

### Vomocytosis in Mucorales: evidence, mechanistic interpretation, and comparison with other pathogens

The statistically significant and reproducible decrease in macrophage phagocytic index from 1.5 to 3 hours for Group 8 at MOI 1 constitutes the first kinetic evidence of vomocytosis in any Mucorales organism, extending the known taxonomy of vomocytosis-capable pathogens from three previously established examples, namely *C. neoformans* (23, 24), *C. albicans* (25), and *A. fumigatus* (26), to a fourth major fungal pathogen group. The key features distinguishing the Group 8 phenotype from the established *Cryptococcus* model are summarised in Table 1.

The inverse dose-dependence observed here, with vomocytosis detected at MOI 1 but absent at MOI 5, contrasts with the *C. neoformans* pattern in which event frequency is positively correlated with intracellular burden (27). This inversion implies an active mechanism: a fungal effector or secreted factor that promotes non-lytic spore release by an as yet uncharacterised mechanism when the macrophage harbours one or a small number of spores, but whose effect is suppressed at higher intracellular loads through the concurrent engagement of NADPH oxidase, LC3-associated phagocytosis, and other intracellular killing mechanisms. The phylogenetic restriction of this phenotype to *Mucor* clade II (*M. heterogamus*, *M. hiemalis*, *M. moelleri*, *M. luteus*) implies conservation of the responsible factor within this clade and its absence or non-expression in the remaining groups tested. Comparative secretomics and surface proteomics between *Mucor* clade I and clade II species represent a rational experimental strategy for identifying this factor.

The higher prevalence of vomocytosis-consistent kinetics in *D. discoideum* relative to MH-S macrophages is attributable to the absence in amoeba of three intracellular retention mechanisms that operate in professional mammalian phagocytes: MARCO scavenger receptor signalling (28), Arp2/3 complex-mediated actin cage formation around pathogen-containing phagosomes, and LC3-associated phagocytosis, which enhances phagolysosomal acidification and fungicidal activity (11). The shared Group 8 vomocytosis signal across both phagocyte types points to a conserved fungal effector of non-host-specific origin. Direct live-cell time-lapse microscopy demonstrating spore extrusion from intact macrophages (23, 24) is required to confirm the vomocytosis interpretation and should be considered the immediate experimental priority.

### The evolutionary training ground hypothesis is not supported at the group level in Mucorales

The complete absence of Spearman rank correlation between macrophage and *D. discoideum* phagocytic indices across all four experimental conditions constitutes a direct experimental test of the evolutionary training ground hypothesis at the order level. The result is unambiguously negative. Macrophage phagocytosis is mediated by Dectin-1, TLR2/4, complement receptor 3, and the mannose receptor, with downstream NADPH oxidase activation, interferon signalling, and adaptive immune crosstalk (11). *D. discoideum* uses lectin-type surface receptors without oxidative burst, cytokine cascade, or adaptive immune coordination (36, 37). These recognition architectures are sufficiently orthogonal that the spore surface features most important for macrophage engagement need not coincide with those most important for amoeba engagement, and the data confirm that they do not.

This conclusion applies to the population-level rank-order prediction. It does not exclude the possibility that individual virulence factors identified through amoeba confrontation experiments retain macrophage-relevant functions, as demonstrated by Itabangi et al. (21) for the *R. microsporus* endosymbiont. The practical consequence is nonetheless clear: *D. discoideum* phagocytic index rankings cannot serve as a validated surrogate for macrophage phagocytic index rankings when assessing the relative innate immune evasion capacity of Mucorales groups.

### Geographic diversity and One Health implications

The dose-dependent difference in macrophage phagocytic index between clinical and environmental isolate groups at MOI 5 but not MOI 1 raises the possibility of cooperative pattern recognition receptor engagement at high spore density, which may be more sensitive to surface ligand density differences between strains with and without mammalian host adaptation. However, since the clinical category comprises only two groups representing distinct genera with different spore dimensions and surface chemistries, this difference cannot be attributed unambiguously to clinical origin per se and is noted as a preliminary observation requiring larger-scale investigation.

The geographic scope of the present collection spans 12 countries across five continents, representing the broadest phagocytosis survey of Mucorales yet assembled. Climate-driven shifts in the geographic range and thermotolerance of Mucorales species are actively reshaping the epidemiology of mucormycosis (38). This dataset establishes a quantitative baseline against which future changes in macrophage interaction phenotype associated with geographic range expansion can be assessed.

### Limitations

Several limitations of this study require acknowledgement. First, phagocytic index measures the proportion of macrophages containing at least one internalised spore, providing no information on intracellular killing efficiency, phagolysosomal maturation, or spore germination fate. Colony-forming unit survival assays and live-cell viability imaging are required to complete the functional characterisation. Second, MH-S cells differ from primary alveolar macrophages in cytokine responsiveness and activation state; validation in primary murine bone marrow-derived macrophages and primary human alveolar macrophages is warranted. Third, Groups 1 and 4 were not included in the *D. discoideum* confrontation assay, limiting the completeness of the cross-system comparison. Fourth, vomocytosis was inferred from kinetic phagocytic index trajectories; direct confirmation by live-cell time-lapse fluorescence microscopy remains the methodological gold standard (23, 24) and is required to exclude alternative interpretations. Fifth, the molecular identity of the Group 8 vomocytosis effector is unknown. Sixth, the macrophage and *D. discoideum* assays were conducted under different physicochemical conditions: 37°C in serum-supplemented RPMI-1640 versus 22°C in Sörensen phosphate buffer. These conditions reflect the natural physiological environments of each cell type, but the contribution of temperature and medium differences to the observed divergence in phagocytic hierarchies cannot be formally excluded.

## CONCLUSIONS

Working through 62 Mucorales strains from nine phylogenetic groups and 12 countries, this study establishes that macrophage phagocytic susceptibility across the order is a reproducible, condition-stable, group-specific phenotype. *Absidia*–*Cunninghamella* species show the lowest macrophage phagocytic indices; *Gilbertella*–*Mycotypha* species show the highest. Group-level mean spore area correlates negatively with macrophage phagocytic index, confirming physical spore dimensions as a virulence-relevant morphometric parameter specific to the mammalian immune context.

The complete absence of rank-order correlation between macrophage and *D. discoideum* phagocytic hierarchies, replicated across all four experimental conditions, demonstrates that the two phagocyte systems engage Mucorales spore surfaces through mechanistically incompatible recognition pathways. Protozoan phagocytic index data cannot serve as a proxy for macrophage phagocytic outcome when ranking Mucorales groups, with direct implications for how amoeba-based Mucorales virulence models should be interpreted.

Kinetic evidence of vomocytosis in *Mucor* clade II organisms, observed in both macrophages and *D. discoideum* with an inverse dose-dependence relative to *Cryptococcus*, points to a conserved active fungal effector. Identification of that effector, direct confirmation of vomocytosis by live-cell imaging, and evaluation of its contribution to virulence in animal models represent the immediate experimental priorities arising from this work.

## Supporting information

Supplemetal tables for measuring spore size

## ACKNOWLEDGEMENTS

We also thank for financial support by the German Egyptian Research Long-term Scholarship (GERLS) Program 2014 (57030312 for M.I.A.H.) coordinated by the German Academic Exchange Service (DAAD) in an initial stage of the study. The authors gratefully acknowledge the support by the Deutsche Forschungsgemeinschaft (DFG) project number 210879364 during the 2^nd^ funding period of the CRC/Transregio124 FungiNet (Projects A6 – KV) in a final phase of the study. We thank the University of Jena and the Leibniz Institute for Natural Product Research and infection Biology (Leibniz-HKI) Jena for excellent working conditions and providing the laboratory facilities during the course of the project. We express our gratitude for curators and the team of the Westerdijk Fungal Biodiversity Institute Utrecht, The Netherlands (formerly: CBS-KNAW); FSU Jena, Germany; SZMC Szeged, Hungary; AUMC, Assiut, Egypt; NRRL Peoria, IL, USA; and MUFS culture collections for providing fungal strains. Statistical analyses and figure generation were performed using Python 3.11 with SciPy, scikit-learn, pandas, and Matplotlib. Language editing assistance was provided by Claude (Anthropic, San Francisco, CA). The scientific content, experimental design, data analysis, and all intellectual contributions are entirely the work of the authors.

## Data Availability

Raw phagocytosis data, corrected strain metadata, and morphometric summary tables are deposited in Zenodo (DOI: to be assigned prior to publication). GenBank accession numbers for ITS and LSU sequences are provided in Supplementary Table S1.

## Author Contributions

M.I.A.H. conceived and designed the study, performed phagocytosis assays and statistical analyses, generated all figures, and wrote the manuscript. Á.C. provided strains, performed phylogenetic analyses, and curated sequence data. K.V. supervised the project, provided resources, and critically revised the manuscript. All authors read and approved the final version.

## Conflicts of Interest

The authors declare no conflicts of interest.

## Funding

This work was supported by the Leibniz Institute for Natural Product Research and Infection Biology – Hans Knöll Institute (Leibniz-HKI), Jena, Germany and the Friedrich Schiller University of Jena and in a final stage of this study also by the Deutsche Forschungsgemeinschaft (DFG) project number 210879364 during the 2^nd^ funding period of the CRC/Transregio124 FungiNet. M.I.A.H. was supported by a DAAD German–Egyptian Long-Term Scholarship (GERLS).

---

**Figure S2.**
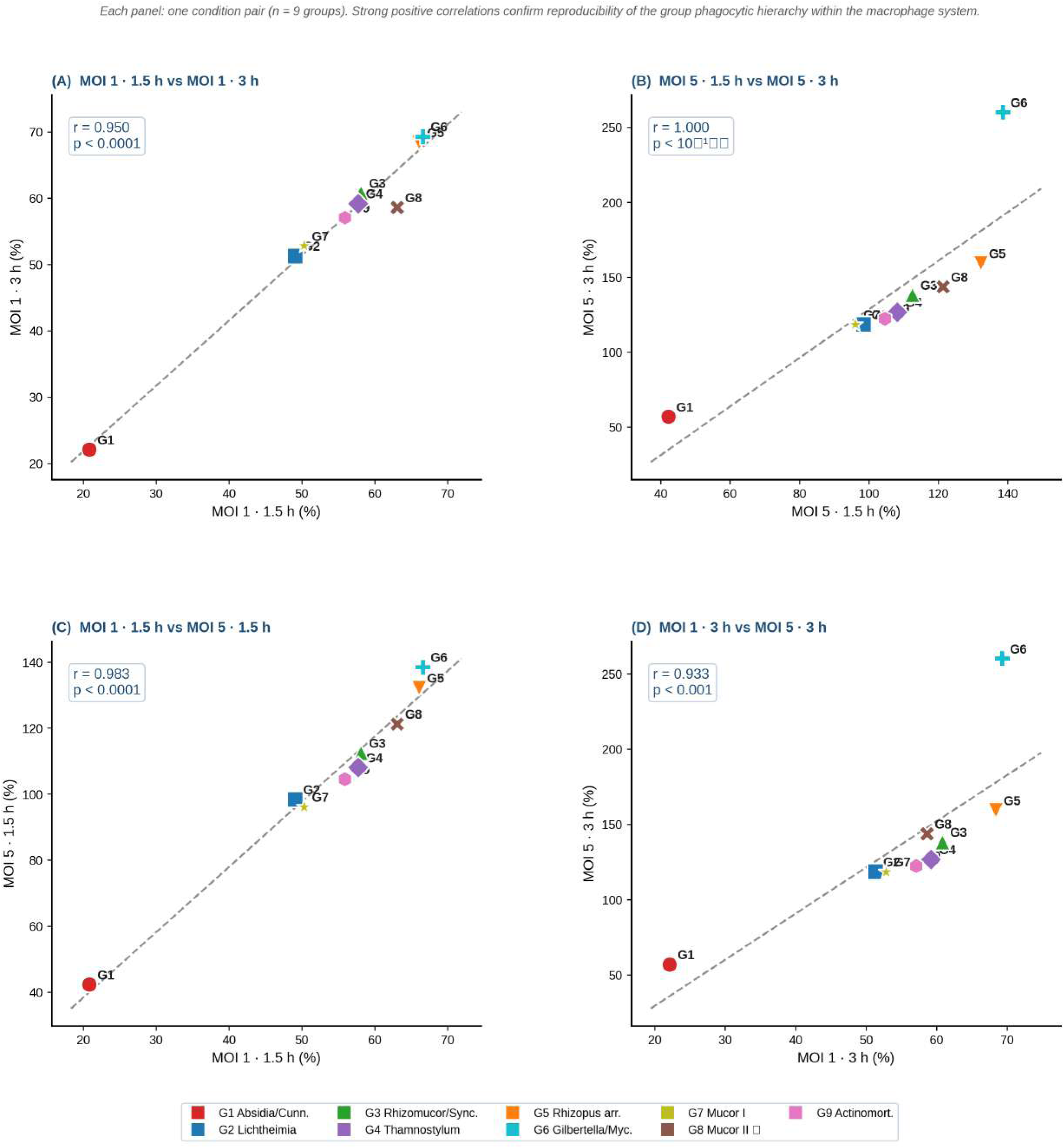
MH-S Macrophage Pl – Internal Spearman Correlations.

**Figure S3.**
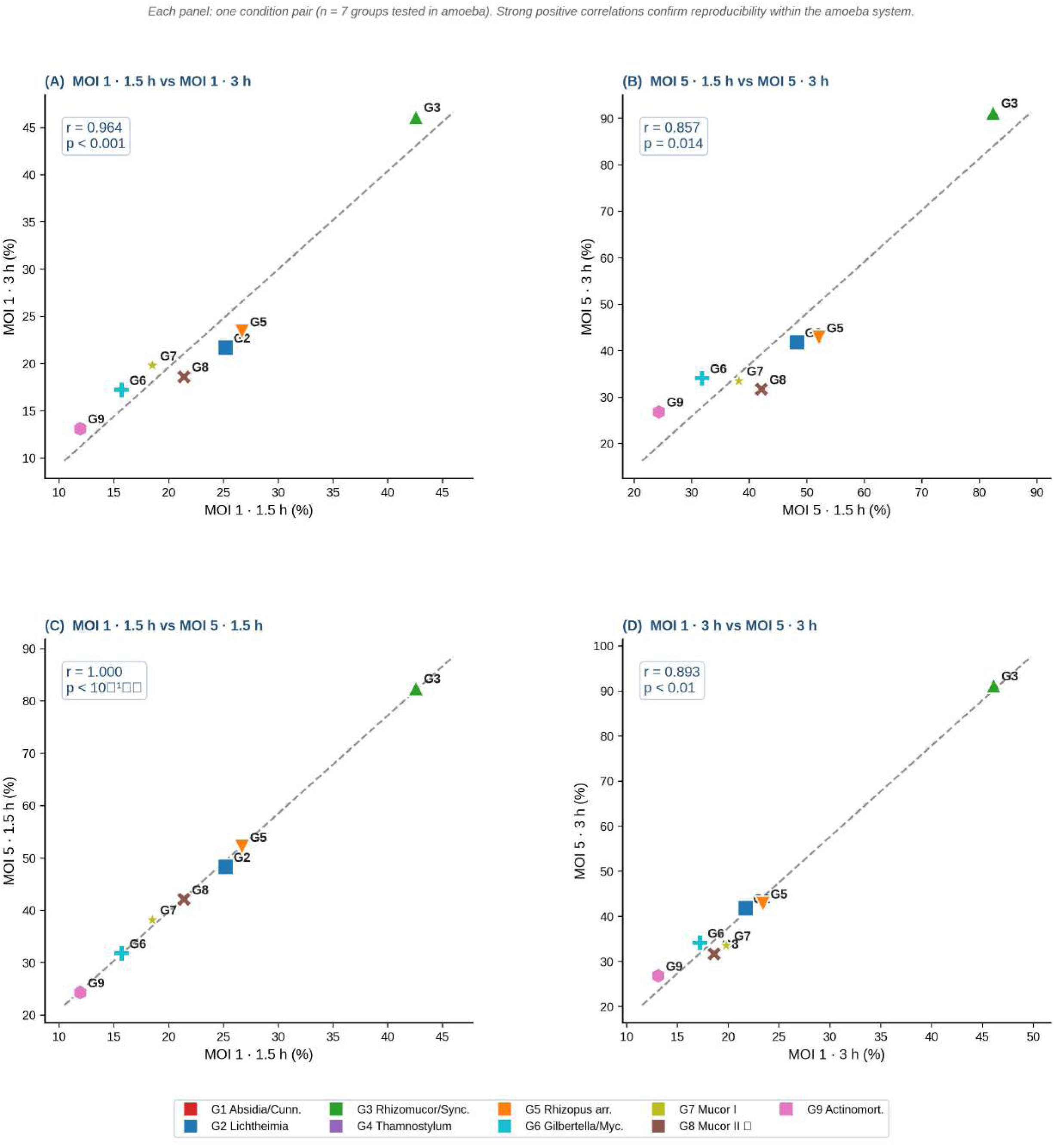
D. discoideum Pl – Internal Spearman Correlations.

**Figure S5.**
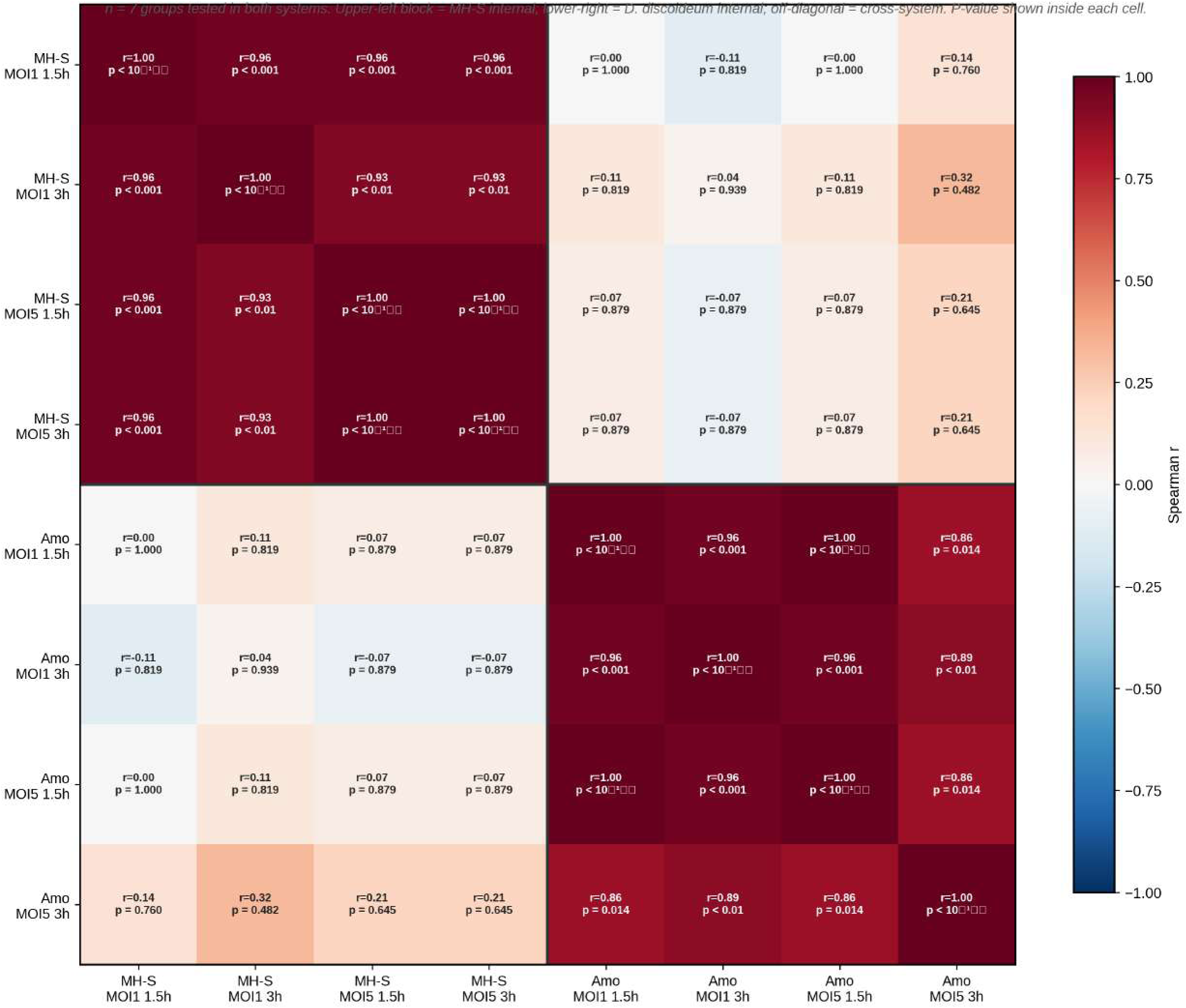
Spearman Rank Correlation Matrix – All Conditions.

## Notes

### Competing Interest Statement

The authors have declared no competing interest.

