## Supplementary material for "Interaction Patterns of Mucormycotina Fungi with Professional Phagocytes and Amoebae: A Comparative Phylogenetic and Geographic Survey on Clinical and Environmental Species": Supplemetal tables for measuring spore size

| Name of species | International number | National number | Area. | Perimeter | Aspect ratio | Solidity |
|---|---|---|---|---|---|---|
| **Group 1** | | | | | | |
| *Absidia* sp1 | FSU:012130 | AUMC-982 | 9.66 ± 2.47 | 11.7 ± 1.50 | 1.21 ± 0.156 | 0.955 ± 0.0167 |
| | FSU:012129 | AUMC-1 | 43.4 ± 82.8 | 19.8 ± 15.3 | 1.33 ± 0.224 | 0.953 ± 0.0248 |
| *Absidia koreana AK* | FSU:012145 | AUMC-6048 | 21.1 ± 7.29 | 17.8 ± 3.45 | 1.60 ± 0.455 | 0.948 ± 0.0246 |
| *Cunninghamella echinulata CE* | FSU:012164 | AUMC-10449 | 56.3 ± 12.6 | 27.2 ± 3.19 | 1.13 ± 0.107 | 0.974 ±0.00560 |
| | FSU:012136 | AUMC-6025 | 42.6 ± 46.3 | 21.0 ± 11.7 | 1.20 ± 0.150 | 0.958 ± 0.0216 |
| | FSU:012140 | AUMC-6030 | 8.34 ± 8.40 | 10.0 ± 4.34 | 1.27 ± 0.177 | 0.956 ± 0.0179 |
| **Group 2** | | | | | | |
| *Lichtheimia corymbifera LC* | FSU:09682 | SZMC 11361 | 462 ± 136 | 78.2 ± 11.2 | 1.24 ± 0.116 | 0.958 ±0.00746 |
| *Lichtheimia hyalospora LH* | FSU:010161 | CBS 102.36 | 573 ± 178 | 86.6 ± 13.0 | 1.14 ± 0.0787 | 0.971 ± 0.0109 |
| | FSU:012133 | AUMC-5707 | 21.9 ± 6.95 | 17.4 ± 2.64 | 1.13 ± 0.0789 | 0.964 ± 0.0120 |
| *Lichtheimia ramosa LR* | FSU:06197 | | 458 ± 195 | 79.3 ± 18.3 | 1.51 ± 0.417 | 0.957 ± 0.0165 |
| | | SZMC 11372 | 372 ± 147 | 72.5 ± 11.9 | 1.60 ± 0.325 | 0.948 ± 0.0155 |
| **Group 3** | | | | | | |
| *Rhizomucor miehei RhM* | CBS 370.71 | SZMC 11028 | 423 ± 137 | 74.3 ± 10.9 | 1.10 ± 0.0655 | 0.959 ± 0.0119 |
| | NRRL 5901 | SZMC 11008 | 371 ± 58.2 | 71.1 ± 5.74 | 1.10 ± 0.0589 | 0.950 ± 0.133 |
| | CBS 360.92 | SZMC 11012 | 350 ± 107 | 67.8 ± 9630 | 1.15 ± 93.4 | 0.953 ± 0.0111 |
| | ETH M4918 | SZMC 11014 | 259 ± 90.3 | 58.3 ± 10.1 | 1.18 ± 0.129 | 0.958 ± 0.0176 |
| *Rhizomucor pusillus RhP* | FSU:012161 | AUMC7966 | 14.0 ± 4.79 | 13.8 ± 2.53 | 1.23 ± 0.239 | 0.964 ± 0.0142 |
| | ETH M4920 | SZMC 11013 | 311 ± 74.2 | 64.0 ± 7.69 | 1.14 ± 0.0803 | 0.953 ± 0.0111 |
| | NRRL 6399 | SZMC 11020 | 291 ± 70.0 | 62.3 ± 7.72 | 1.14 ± 0.106 | 0.949 ± 0.0106 |
| | WRL CN(M) 231 | SZMC 11015 | 416 ± 102 | 74.2 ± 9.18 | 1.16 ± 0.118 | 0.966 ± 0.0103 |
| *Syncephalastrum monosporum SM* | | SZMC 0047 | 669 ± 309 | 93.0 ± 18.1 | 1.11 ± 0.0700 | 0.965 ± 0.0700 |
| *Syncephalastrum racemosum SR* | FSU:012160 | AUMC7965 | 122 ± 66.4 | 39.0 ± 10.6 | 1.13 ± 0.0755 | 0.972 ± 0.00656 |
| | FSU:012148 | AUMC-6115 | 16.1 ± 4.35 | 15.2 ± 2.35 | 1.32 ± 0.354 | 0.954 ± 0.136 |
| **Group 4** | | | | | | |
| *Thamnostylum piriforme TP* | MUFS 053 | SZMC 22673 | 446 ± 126 | 78.0 ± 10.9 | 1.24 ± 0.196 | 0.954 ± 0.0130 |
| **Group 5** | | | | | | |
| *Rhizopus arrhizus RO* | FSU 8743 | SZMC 21290 | 564 ± 146 | 87.1 ± 11.0 | 1.19 ± 0.117 | 0.963 ± 0.0121 |
| | FSU 5857 | SZMC 21291 | 659 ± 179 | 93.2 ± 13.1 | 1.17 ± 0.0901 | 0.966 ±0.00799 |
| | FSU:012152 | AUMC7959 | 27.9 ± 9.39 | 19.4 ± 3.20 | 1.19 ± 0.130 | 0.969 ± 0.192 |
| | RA 99-880 | SZMC 13635 | 144 ± 39.2 | 44.4 ± 6.20 | 1.25 ± 0.168 | 0.943 ± 0.0266 |
| | | SZMC1107 | 440 ± 99.2 | 76.8 ± 8.47 | 1.21 ± 0.128 | 0.961 ±0.00876 |
| | TJM7F2 | SZMC 13611 | 558 ± 122 | 86.3 ± 9.38 | 1.20 ± 0.123 | 0.962 ± 0.00809 |
| | TJM24B | SZMC 11101 | 1080 ± 342 | 120 ± 18.3 | 1.28 ± 0.241 | 0.969 ± 0.0117 |
| *Rhizopus microsporus RM* | | SZMC 21297 | 199 ± 34.9 | 52.1 ± 4.61 | 1.19 ± 0.114 | 0.942 ± 0.0199 |
| | NRRL 2710 | SZMC 13622 | 1520 ± 501 | 142 ± 24.4 | 1.20 ± 0.180 | 0.967 ± 0.0105 |
| | CBS 102.277 | SZMC 13645 | 648 ± 155 | 92.5 ± 11.2 | 1.13 ± 0.147 | 0.965 ±0.00823 |
| **Group 6** | | | | | | |

| | | | | | | |
|---|---|---|---|---|---|---|
| *Gilbertella persicaria GP* | | SZMC 11099 | 644 ± 119 | 93.0 ± 8.64 | 1.16 ± 0.111 | 0.961 ±0.00723 |
| *Mycotypha africana MyA* | CBS 122.64 | SZMC 11069M | 479 ± 153 | 83.1 ± 12.6 | 1.48 ± 0.343 | 0.949 ± 0.0228 |
| | FSU 296 | SZMC 11070M | 87.6 ± 24.2 | 34.5 ± 4.69 | 1.31 ± 0.190 | 0.940 ± 0.0282 |
| *Mycotypha microspora MyM* | - | SZMC 0471 | 157 ± 64.5 | 45.2 ± 8.56 | 1.18 ± 0.118 | 0.948 ± 0.0195 |
| **Group7** | | | | | | |
| *Ellisomyces anomalus EA* | CBS 243.57 | SZMC 23391 | 2420 ± 3280 | 161 ± 90.6 | 1.18 ± 0.142 | 0.966 ± 0.0112 |
| *Mucor circinelloides MC* | FSU:012141 | AUMC-6031 | 21.6 ± 6.11 | 17.1 ± 2.26 | 1.17 ± 0.107 | 0.967 ±0.00888 |
| | FSU:012142 | AUMC-6033 | 19.8 ± 4.93 | 116.6 ± 2.05 | 1.18 ± 0.105 | 0.967 ±0.00815 |
| | FRR 2109 | SZMC 12028 | 677 ± 266 | 95.3 ± 18.3 | 1.36 ± 0.240 | 0.963 ±0.00687 |
| | NRRL 3632 | SZMC 12054 | 935 ± 152 | 112 ± 9.29 | 1.23 ± 0.163 | 0.965 ±0.00533 |
| *Mucor janssenii MJ* | CBS 243.67 | SZMC 12003 | 695 ± 296 | 94.7 ± 18.9 | 1.13 ± 0.0817 | 0.962 ±0.00840 |
| *Mucor lusitanicus ML* | MS12 | SZMC 12082 | 1520 ± 1570 | 135 ± 67.8 | 1.47 ± 0.283 | 0.958 ± 0.0143 |
| | MUFS 029 | SZMC 12069 | 610 ± 235 | 89.7 ± 16.8 | 1.19 ± 0.148 | 0.964 ± 0.0155 |
| | ATCC 1216 | SZMC 12030 | 623 ± 120 | 92.0 ± 8.86 | 1.25 ± 0.184 | 0.962 ± 0.0102 |
| | CBS 277.49 | | 2140 ± 994 | 168 ± 41.4 | 1.37 ± 0.235 | 0.966 ±0.00944 |
| *Mucor plumbeus MP* | MUFS 162 | SZMC 12070 | 1260 ± 398 | 129 ± 20.1 | 1.13 ± 0.0790 | 0.971 ±0.00653 |
| *Mucor racemosus MR* | G423 | SZMC 12039 | 619 ± 546 | 82.5 ± 40.4 | 1.26 ± 0.224 | 0.957 ± 0.0942 |
| | MUFS 055 | SZMC 12067 | 1120 ± 342 | 121 ± 19.0 | 1.16 ± 0.0949 | 0.970 ± 0.122 |
| | MUFS 056 | SZMC 12068 | 882 ± 268 | 108 ± 17.6 | 1.23 ± 0.102 | 0.967 ±0.00733 |
| *Mucor pseudolusitanicus MPs* | FSU:12135 | AUMC-6022 | 19.7 ± 6.12 | 17.0 ± 2.71 | 1.47 ± 0.317 | 0.958 ± 0.0139 |
| **Group 8** | | | | | | |
| *Actinomucor elegans AC* | FSU: 012134 | AUMC-6020 | 32.4 ± 7.85 | 20.9 ± 2.39 | 1.12 ± 0.0606 | 0.970 ± 0.0111 |
| | FSU: 012144 | AUMC-6040 | 7.50 ± 7.73 | 9.20 ± 4.37 | 1.19 ± 0.138 | 0.960 ± 0.0230 |
| | | SZMC 00250 | 1440 ± 742 | 136 ± 34.6 | 1.19 ± 0.158 | 0.965 ± 0.0117 |
| | FSU:012146 | AUMC-6050 | 26.3 ± 8.30 | 18.6 ± 2.92 | 1.19 ± 0.141 | 0.973 ± 0.136 |
| *Mucor heterogamus MHe* | NRRL2663, ATCC36727 | SZMC 11073 | 383 ± 95.6 | 71.1 ± 9.15 | 1.16 ± 0.0963 | 0.867 ± 0.0701 |
| *Mucor hiemalis MHi* | NRRL 3624 | SZMC 12056 | 960 ± 402 | 112 ± 24.6 | 1.32 ± 0.142 | 0.967 ±0.00882 |
| *Mucor moelleri MM* | FSU:012131 | AUMC-3674 | 14.9 ± 4.95 | 14.9 ± 2.53 | 1.46 ± 0.451 | 0.954 ± 0.136 |
| **Group 9** | | | | | | |
| *Actinomortierella wolfii AW* | CBS 209.69 | SZMC 11245 | 494 ± 139 | 80.8 ± 11.2 | 1.18 ± 0.107 | 0.957 ±0.00725 |

| Name of species | International number | National number | Area. | Perimeter | Aspect ratio | Solidity |
|---|---|---|---|---|---|---|
| **Group 1** | | | | | | |
| *Absidia* sp1 | FSU:012130 | AUMC-982 | 11.5 ± 3.06 | 13.7 ± 2.05 | 1.25 ± 0.157 | 0.938 ± 0.188 |
| | FSU:012129 | AUMC-1 | 75.7 ± 92.5 | 27.8 ±17.4 | 1.27 ± 0.178 | 0.963 ± 0.016 |
| *Absidia koreana AK* | FSU:012145 | AUMC-6048 | 27.6 ± 19.9 | 21.4 ± 7.79 | 1.68 ± 0.590 | **0.920 ± 0.0413** |
| *Cunninghamella echinulata CE* | FSU:012164 | AUMC-10449 | 90.0 ± 28.0 | 34.7 ± 5.72 | 1.13 ± 0.0895 | 0.973 ± 0.00997 |
| | FSU:012136 | AUMC-6025 | 141 ± 87.5 | 44.2 ± 13.5 | 1.20 ± 0.131 | 0.960 ± 0.207 |
| | FSU:012140 | AUMC-6030 | 40.0 ± 21.6 | 24.2 ± 6.76 | 1.28 ± 0.183 | 0.930 ± 0.0379 |
| **Group 2** | | | | | | |
| *Lichtheimia corymbifera LC* | FSU:09682 | SZMC 11361 | 662 ±226,61 | 93.6 ± 15.9778 | 1.18 ± 0.107 | 0.959 ± 0.01008 |
| *Lichtheimia hyalospora LH* | FSU:010161 | CBS 102.36 | 599 ± 197 | 88.7 ± 13.8 | 1.12 ± 0.0723 | 0.963 ± 0.00848 |
| | FSU:012133 | AUMC-5707 | 23.4 ± 6.15 | 18.9 ± 2.50 | 1.21 ± 0.155 | 0.951 ± 0.136 |
| *Lichtheimia ramosa LR* | FSU:06197 | | 461 ± 173 | 79.2 ± 14.5 | 1.34 ± 0.195 | 0.956 ± 0.0152 |
| | | SZMC 11372 | 429 ± 118 | 77.3 ± 10.8 | 1.48 ± 0.252 | 0.959 ± 0.0129 |
| **Group 3** | | | | | | |
| *Rhizomucor miehei RhM* | CBS 370.71 | SZMC 11028 | 483 ± 182 | 79.1 ± 14.5 | 1.16 ± 0.107 | 0.964 ± 0.0101 |
| | NRRL 5901 | SZMC 11008 | 448 ± 110 | 77.0 ± 9.07 | 1.13 ± 0.0897 | 0.960 ± 0.00869 |
| | CBS 360.92 | SZMC 11012 | 596 ± 213 | 88.9 ± 13.3 | 1.15 ± 0.0801 | 0.961 ± 0.00961 |
| | ETH M4918 | SZMC 11014 | 310 ± 65.9 | 64.3 ± 6.99 | 1.18 ± 0.123 | 0.955 ± 0.00941 |
| *Rhizomucor pusillus RhP* | FSU:012161 | AUMC7966 | 14.7 ± 6.19 | 14.3 ± 2.60 | 1.19 ± 0.131 | 0.965 ± 0.0105 |
| | ETH M4920 | SZMC 11013 | 435 ± 127 | 75.4 ± 10.8 | 1.11 ± 0.0749 | 0.959 ± 0.185 |
| | NRRL 6399 | SZMC 11020 | 372 ± 129 | 70.1 ± 12.0 | 1.12 ± 0.0807 | 0.956 ± 0.0085 |
| | WRL CN(M) 231 | SZMC 11015 | 438 ± 132 | 76.1 ± 10.4 | 1.16 ± 0.0992 | 0.964 ± 0.00768 |
| *Syncephalastrum monosporum SM* | | SZMC 0047 | 999 ± 337 | 114 ± 19.7 | 1.10 ± 0.0769 | 0.969 ± 0.00872 |
| *Syncephalastrum racemosum SR* | FSU:012160 | AUMC7965 | 159 ± 87.6 | 44.1 ± 12.3 | 1.09 ± 0.0480 | 0.975 ± 0.00523 |
| | FSU:012148 | AUMC-6115 | 17.0 ± 7.10 | 15.5 ± 2.89 | 1.16 ± 0.114 | 0.959 ± 0.136 |
| **Group 4** | | | | | | |
| *Thamnostylum piriforme TP* | MUFS 053 | SZMC 22673 | 512 ± 211 | 81.7 ± 16.9 | 1.15 ± 0.127 | 0.960 ± 0.0118 |
| **Group 5** | | | | | | |
| *Rhizopus arrhizus RO* | FSU 8743 | SZMC 21290 | 867 ± 353 | 106 ± 20.5 | 1.21 ± 0.127 | 0.969 ± 0.00718 |
| | FSU 5857 | SZMC 21291 | 922 ± 418 | 108 ± 25.1 | 1.12 ± 0.0789 | 0.965 ± 0.00982 |
| | FSU:012152 | AUMC7959 | 28.6 ± 13.6 | 19.8 ± 4.66 | 1.23 ± 0.181 | 0.962 ± 0.136 |
| | RA 99-880 | SZMC 13635 | 623 ± 161 | 90.7 ± 12.3 | 1.15 ± 0.114 | 0.963 ± 0.00708 |
| | | SZMC 11027 | 743 ± 259 | 99.0 ± 15.5 | 1.18 ± 0.127 | 0.965 ± 0.00800 |
| | TJM7F2 | SZMC 13611 | 520 ± 107 | 83.2 ± 8.37 | 1.17 ± 0.0990 | 0.961 ± 0.00669 |
| | TJM24B | SZMC 11101 | 1110 ± 403 | 122 ± 20.5 | 1.22 ± 0.135 | 0.964 ± 0.124 |
| *Rhizopus microsporus RM* | | SZMC 21297 | 589 ± 139 | 88.1 ± 9.97 | 1.11 ± 0.0631 | 0.960 ± 0.00668 |
| | NRRL 2710 | SZMC 13622 | 1580 ± 473 | 145 ± 21.9 | 1.19 ± 0.126 | 0.966 ± 0.00852 |
| | CBS 102.277 | SZMC 13645 | 650 ± 193 | 92.1 ± 13.1 | 1.11 ± 0.0786 | 0.963 ± 0.00871 |

| | | | | | | |
|---|---|---|---|---|---|---|
| **Group 6** | | | | | | |
| *Gilbertella persicaria GP* | | SZMC 11099 | 744 ± 212 | 99.8 ± 14.4 | 1.15 ± 0.102 | 0.961 ± 0.130 |
| *Mycotypha africana MyA* | CBS 122.64 | SZMC11069 M | 455 ± 115 | 81.8 ± 10.6 | 1.54 ± 0.336 | 0.950 ± 0.0148 |
| | FSU 296 | SZMC11070 M | 409±135 | 76.2±12.4 | 1.55 ± 0.278 | 0.953 ± 0.0136 |
| *Mycotypha microspora MyM* | - | SZMC 0471 | 327 ± 108 | 66.1 ± 10.7 | 1.21 ± 0.16 | 0.954 ± 0.0121 |
| **Group7** | | | | | | |
| *Ellisomyces anomalus EA* | CBS 243.57 | SZMC 23391 | 3080 ± 3490 | 178 ± 104 | 1.20 ± 0.172 | 0.967 ± 0.0114 |
| *Mucor circinelloides MC* | FSU:012141 | AUMC-6031 | 29.5 ± 11.4 | 22.3 ± 4.88 | 1.28 ± 0.243 | 0.94 ± 0.226 |
| | FSU:012142 | AUMC-6033 | 27.5 ± 13.2 | 21.4 ± 5.16 | 5.20 ± 1.28 | 0.799 ± 0.0885 |
| | FRR 2109 | SZMC 12028 | 695 ± 226 | 95.3 ± 14.2 | 1.14 ± 0.106 | 0.965 ± 0.00652 |
| | NRRL 3632 | SZMC 12054 | 1100 ± 370 | 120 ± 18.4 | 1.20 ± 0.113 | 0.968 ± 0.00805 |
| *Mucor janssenii MJ* | CBS 243.67 | SZMC 12003 | 846 ± 260 | 105 ± 15.9 | 1.11 ± 0.0593 | 0.966 ± 0.00779 |
| *Mucor lusitanicus ML* | MS12 | SZMC12082 | 3290 ± 2200 | 203 ± 76.3 | 1.27 ± 0.229 | 0.967 ± 0.0251 |
| | MUFS 029 | SZMC 12069 | 755 ± 218 | 99.7 ± 13.8 | 1.18 ± 0.127 | 0.968 ± 0.00758 |
| | ATCC 1216 | SZMC 12030 | 734 ± 220 | 98.0 ± 14.2 | 1.13 ± 0.0764 | 0.967 ± 0.00618 |
| | CBS 277.49 | | 2330 ± 2200 | 161 ± 79.1 | 1.30 ± 0.243 | 0.970 ± 0.00924 |
| *Mucor plumbeus MP* | MUFS 162 | SZMC 12070 | 1260 ± 396 | 128 ± 19.8 | 1.11 ± 0.0976 | 0.971 ± 0.00721 |
| *Mucor racemosus MR* | G423 | SZMC 12039 | 856 ± 334 | 105 ± 22.5 | 1.17 ± 0.0905 | 0.965 ± 0.00792 |
| | MUFS 055 | SZMC 12067 | 1270 ± 435 | 128 ± 24.1 | 1.15 ± 0.0969 | 0.966 ± 0.00764 |
| | MUFS 056 | SZMC 12068 | 1070 ± 326 | 119 ± 18.2 | 1.15 ± 0.0913 | 0.966 ± 0.00689 |
| *Mucor pseudolusitanicus MPs* | FSU:12135 | AUMC-6022 | 25.9 ± 13.4 | 19.9 ± 5.08 | 1.47 ± 0.325 | 0.947 ± 0.1381 |
| **Group 8** | | | | | | |
| *Actinomucor elegans AC* | FSU: 012134 | AUMC-6020 | 50.4 ± 16.0 | 27.1 ± 4.00 | 1.24 ± 0.134 | 0.960 ± 0.0173 |
| | FSU: 012144 | AUMC-6040 | 32.8 ± 18.7 | 23.2 ± 6.78 | 1.25 ± 0.133 | 0.921 ± 0.150 |
| | | SZMC 00250 | 1730 ± 655 | 150 ± 28.7 | 1.12 ± 0.0842 | 0.970 ± 0.00676 |
| | FSU:012146 | AUMC-6050 | 25.0 ± 9.29 | 19.3 ± 3.40 | 1.24 ± 0.160 | 0.954 ± 0.226 |
| *Mucor heterogamus MHe* | NRRL 2663, ATCC 36727 | SZMC 11073 | 364 ± 88.2 | 69.0 ± 8.71 | 1.12 ± 0.0751 | 0.961 ± 0.00812 |
| *Mucor hiemalis MHi* | NRRL 3624 | SZMC 12056 | 948 ± 351 | 112 ± 21.5 | 1.29 ± 0.184 | 0.967 ± 0.00954 |
| *Mucor moelleri MM* | FSU:012131 | AUMC-3674 | 15.8 ± 6.16 | 15.8 ± 3.03 | 1.35 ± 0.351 | 0.944 ± 0.0332 |
| **Group 9** | | | | | | |
| *Actinomortierella wolfii AW* | CBS 209.69 | SZMC 11245 | 676 ± 249 | 94.0 ± 15.9 | 1.14 ± 0.0841 | 0.963 ± 0.00623 |

| Groups | Area | | Perimeter | | Aspect Ratio | | Solidity | |
|---|---|---|---|---|---|---|---|---|
| | 1.5 h | 3h | 1.5 h | 3h | 1.5 h | 3 h | 1.5 h | 3 h |
| Group 1 | 30.2 ± 26.6 | 67.2 ± 42.1 | 17.9 ± 6.58 | 27.7 ± 8.87 | 1.29 ±0.212 | 1.30 ±0.221 | 0.957 ±0.0185 | 0.947 ± 0.0834 |
| Group 2 | 377 ± 133 | 435 ± 144 | 66.8 ± 11.4 | 71.5 ± 11.5 | 1.26 ±0.173 | 1.27 ±0.156 | 0.960 ±0.0125 | 0.958 ± 0.0365 |
| Group 3 | 295 ± 93.0 | 388 ± 127 | 57.5±8.83 | 65.3 ± 10.4 | 1.16 ±0.0861 | 1.14±0.093 | 0.958 ±0.0393 | 0.962 ± 0.0363 |
| Group 4 | 446 ± 126 | 512 ± 211 | 78.0 ± 10.9 | 81.7 ± 16.9 | 1.24 ± 0.196 | 1.15 ±0.127 | 0.954 ±0.0130 | 0.960 ± 0.0118 |
| Group 5 | 584 ± 163 | 831 ± 248 | 81.4 ± 11.0 | 105 ± 14.9 | 1.21 ±0.143 | 1.16 ±0.105 | 0.960±0.033 | 0.964 ± 0.0323 |
| Group 6 | 342 ± 90.2 | 484 ± 143 | 64 ± 8.62 | 81 ± 12 | 1.32 ±0.215 | 1.43 ±0.258 | 0.950 ±0.0194 | 0.952 ± 0.0135 |
| Group 7 | 904 ± 566 | 1160 ± 714 | 95.9 ± 25.1 | 113 ± 29.0 | 1.25 ±0.167 | 1.47 ±0.223 | 0.965 ±0.0235 | 0.953 ± 0.0376 |
| Group 8 | 409 ± 181 | 452 ± 163 | 61.3 ± 13.0 | 59.5 ± 10.9 | 1.23 ± 0.17 | 1.23 ± 0.16 | 0.950 ±0.0435 | 0.954 ± 0.0644 |
| Group 9 | 494 ± 139 | 676 ± 249 | 80.8 ± 11.2 | 94.0 ± 15.9 | 1.18 ±0.107 | 1.14 ±0.0841 | 0.957±0.0072 | 0.963 ± 0.00623 |
